# Revolutionising the design and analysis of protein engineering experiments using fractional factorial design

**DOI:** 10.1101/298273

**Authors:** Steven P. D. Harborne, Duncan Wotherspoon, Jessica Michie, Alasdair McComb, Tommi Kotila, Steven G. Gilmour, Adrian Goldman

**Affiliations:** Astbury Centre for Structural Biology, School of Biomedical Sciences, Faculty of Biological Sciences, University of Leeds, LS2 9JT, UK; Division of Biochemistry, Biological and Environmental Sciences, University of Helsinki, Finland; Department of Mathematics, King’s College London, Strand, London, WC2R 2LS, UK

**Keywords:** Incomplete factorial design, X-ray crystallography, mutagenesis, protein engineering, purification

## Abstract

Protein engineering is one of the foundations of biotechnology, used to increase protein stability, re-assign the catalytic properties of enzymes or increase the interaction affinity between antibody and target. To date, strategies for protein engineering have focussed on systematic, random or computational methods for introducing new mutations. Here, we introduce the statistical approach of fractional factorial design as a convenient and powerful tool for the design and analysis of protein mutations, allowing sampling of a large mutational space whilst minimising the tests to be done. Our test case is the integral membrane protein, Acridine resistance subunit B (AcrB), part of the AcrAB-TolC multi-protein complex, a multi-drug efflux pump of Gram-negative bacteria. *E. coli* AcrB is naturally histidine-rich, meaning that it is a common contaminant in the purification of recombinantly expressed, histidine-tagged membrane proteins. Coupled with the ability of AcrB to crystallise from picogram quantities causing false positives in 2-D and 3-D crystallisation screening, AcrB contamination represents a significant hindrance to the determination of new membrane protein structures. Here, we demonstrate the use of fractional factorial design for protein engineering, identifying the most important residues involved in the interaction between AcrB and nickel resin. We demonstrate that a combination of spatially close, but sequentially distant histidine residues are important for nickel binding, which were different from those predicted *a priori*. Fractional factorial methodology has the ability to decrease the time and material costs associated with protein engineering whilst expanding the depth of mutational space explored; a revolutionary concept.

**Significance statement:** Protein engineering is important for the production of enzymes for bio-manufacturing, stabilised protein for research and production of therapeutic antibodies against human diseases. Here, we introduce a statistical method that can reduce the time and cost required to perform protein engineering. We validate our approach experimentally using the multi-drug efflux pump AcrB, a target for understanding drug-resistance in pathogenic bacteria, but also a persistent contaminant in the purification of membrane proteins from *E. coli*. This provides a general method for increasing the efficiency of protein engineering.

## Introduction

Protein engineering is an extremely useful tool in protein biotechnology for applications such as protein stabilisation, re-assigning the catalytic properties of enzymes or increasing the interaction affinity between antibody and target. However, a problem arises in the fact that for a protein of N residues, the number of possible sequences is 20^N^. Therefore, for a 300-residue protein the number of possible sequences is 20^300^ - effectively infinite possibilities. Even scaling this back to consider just a small subset of positions for mutation provides a colossal number of potential mutations (the mutation space), which remains a major problem for understanding protein folding and improving protein function for biotechnological purposes. To date, strategies for protein engineering have focussed on scanning (1), semi-systematic (2, 3), random (4), directed evolution (5) or computational methods (6–8) for introducing new mutations.

Scanning mutagenesis has been particularly popular for the stabilisation of GPCRs and has had highly successful outcomes for the structural elucidation of this extremely important class of membrane protein (9). However, the process by which mutations are made and selected is an expensive and labour-intensive process due to the fact that every amino acid position must be mutated individually and tested for changes to protein behaviour (*e.g*. thermostability), and therefore this approach has been somewhat exclusive to industry. Furthermore, scanning techniques are limited to finding single positions at a time, and provide no information about additive effects of combined mutations. Instead, amino acid positions initially identified by scanning have traditionally been combined in a semi-systematic way (2), and from previous evidence it is clear that combining single mutations together rarely provides a straight-forward additive effect (2).

Alternatively, mutagenesis can be performed randomly using techniques such as error prone PCR (10) or mutator strains of *E. coli* (11). These random methods can be used for directed evolution by multiple iterations of random mutagenesis followed by screening. However, these methods requires the use of rapid, robust and high-throughput assays for evaluating mutational outcomes (for example levels of GFP fluorescence (5)) and a method to link improvement in function to the sequence that gave rise to it (for example cell sorting (5) or phage display (12)). However, not all strategies are amenable to these approaches, as improvements in function may require complex assays to ascertain. Furthermore, approaches that rely on error-prone PCR are limited due to several compounding factors. Primarily, certain base-changes are more common than others (10), for example A for T substitutions are more common than C for G substitutions (10). Secondly, a single base-pair change to a codon is insufficient for one amino acid to be changed into all other amino acids, for example, with a single base-pair change, alanine can be mutated to valine, threonine, proline, serine, aspartate, glutamate or glycine, but not to anything else. Under conventional error-prone PCR methods, a double base-pair change in a single codon is statistically unlikely; therefore, the kinds of changes that can be made to amino acid sequence using error-prone PCR are biased and limited.

Computational methods for predicting and designing advantageous changes to protein sequence are in their infancy (6). There have been several notable examples of where this approach has been successful (7, 8), but often relies on pre-existing structural information (which is not always available) and high-level thermodynamic calculations. Alternatively, deep sequencing information has been exploited, for example the availability of homologs from thermophilic or thermotolerant organisms has helped to successfully predict mutations for thermostabilisation of certain membrane proteins (13). However, not all proteins of interest will have thermostable homologs in nature.

Here, we intend to introduce a statistical method that will be widely applicable to protein engineering, and pose some significant advantages over other approaches. Our key observation is that each residue on average interacts with just three or four others, and most of the effects of mutating a residue will be due to these local interactions. We can sample this space efficiently by devising a mutation strategy that focuses only on minimal changes. Such a strategy is called a fractional factorial design. A full factorial design would be one in which there are a number of ‘factors’ to be tested (i.e. interesting residue positions to mutate) each of which has a number of discrete ‘levels’ (i.e. mutated or not mutated, or mutated to one of 20 amino acids) and every combination of these levels across all factors would be tested. A fractional factorial design consists of a carefully selected subset of the combinations available in a full factorial design, chosen to exploit the sparsity-of-effects principal and reveal the most important information about the system being studied. Fractional factorial approaches have become an important part of the statistical toolkit in mechanical engineering and pharmaceutical science, and we intend to apply it to our case of protein engineering.

Fractional factorial approaches have been tried on occasion in protein science: Carter and Carter in 1979 (14) proposed their use for protein crystallisation, but this approach has been completely superseded by knowledge-based “sparse-matrix” screens (15). Recently, the fractional factorial approach was used to optimise protein expression. The factors included different fusion tags, strains and growth media (16), allowing a more efficient approach to optimising the conditions than a full factorial design, similar to much earlier work on process optimisation (17). However, none of this work has focused directly on optimising protein sequence.

To demonstrate the ability of fractional factorial design as a useful tool in protein engineering we have selected the test case of Acridine resistance subunit B (AcrB) from *E. coli*, which is part of the AcrAB-TolC multi-protein complex, a multi-drug efflux pump of Gram-negative bacteria. Export proteins such as AcrB have emerged as important players for the clinical treatment of infectious disease due to the fact that these proteins confer resistance in Gram-negative bacteria (such as *Salmonella*) to antibiotics, detergents and cationic dyes among others. Aside from its importance as a target for understanding drug-resistance in pathogenic bacteria, AcrB also has considerable implications in the field structural biology as *E. coli* AcrB has often been reported as a contaminant in membrane protein preparations prior to X-ray crystallography (18–20); it is naturally histidine-rich and therefore readily binds to charged nickel resins (19). As little as picogram quantities of contaminating AcrB can lead to the formation of characteristic rhombohedral crystals (20). Highlighting this issue is a report that of 17 integral membrane protein candidates from *Helicobacter pylori* over expressed in *E. coli*, 45% of crystal hits were discovered to be AcrB crystals (20).

The routine contamination of AcrB is in part due to the fact that levels of AcrAB transcription are inversely proportional to the bacterial rate of growth (21). AcrB expression is therefore greatest in the late stationary phase of growth, as induced by standard laboratory over-expression methods. Furthermore, increasing the stringency of purification steps proven effective in the elimination of other contaminants such as succinate dehydrogenase (20), fails for AcrB due to its particularly high affinity for nickel, thus, making it very difficult to remove by conventional means, resulting in its co-purification alongside his-tagged proteins of interest.

Deleting the histadine rich C-terminus of AcrB has not been successful, and *E. coli* strains with inactive AcrB (*ΔAcrB*) tend to be more sensitive to antibiotics (22) a serious concern for the use of over-expression systems. Therefore, a better approach is to introduce the minimal number of changes required to reduce the affinity of *E. coli* AcrB to nickel sepharose resin to produce functional AcrB with reduced affinity for nickel that can replace wild-type AcrB in *E. coli* expression strains. Furthermore, success in this goal will demonstrate the validity and strengths of fractional factorial design as a valuable tool for protein engineering.

## Results

*E. coli* AcrB has eleven histidine residues per protomer (33 across the trimer), of which seven (H505, H525, H526, H1042, H1044, H1048 and H1049) are clustered on the cytoplasmic proximal face (Fig. 1). Due to their proximity to one another and position on the surface of the protein these seven histidine residues were selected as likely candidates for the innate affinity of AcrB for nickel. To investigate the possible contribution of these residues to nickel binding, we used a fractional factorial design to distinguish primary effects of individual mutations (*main effects*) from pairwise effects of two residues acting together synergistically (*two-way effects*) (Table 1).

**Figure 1.**
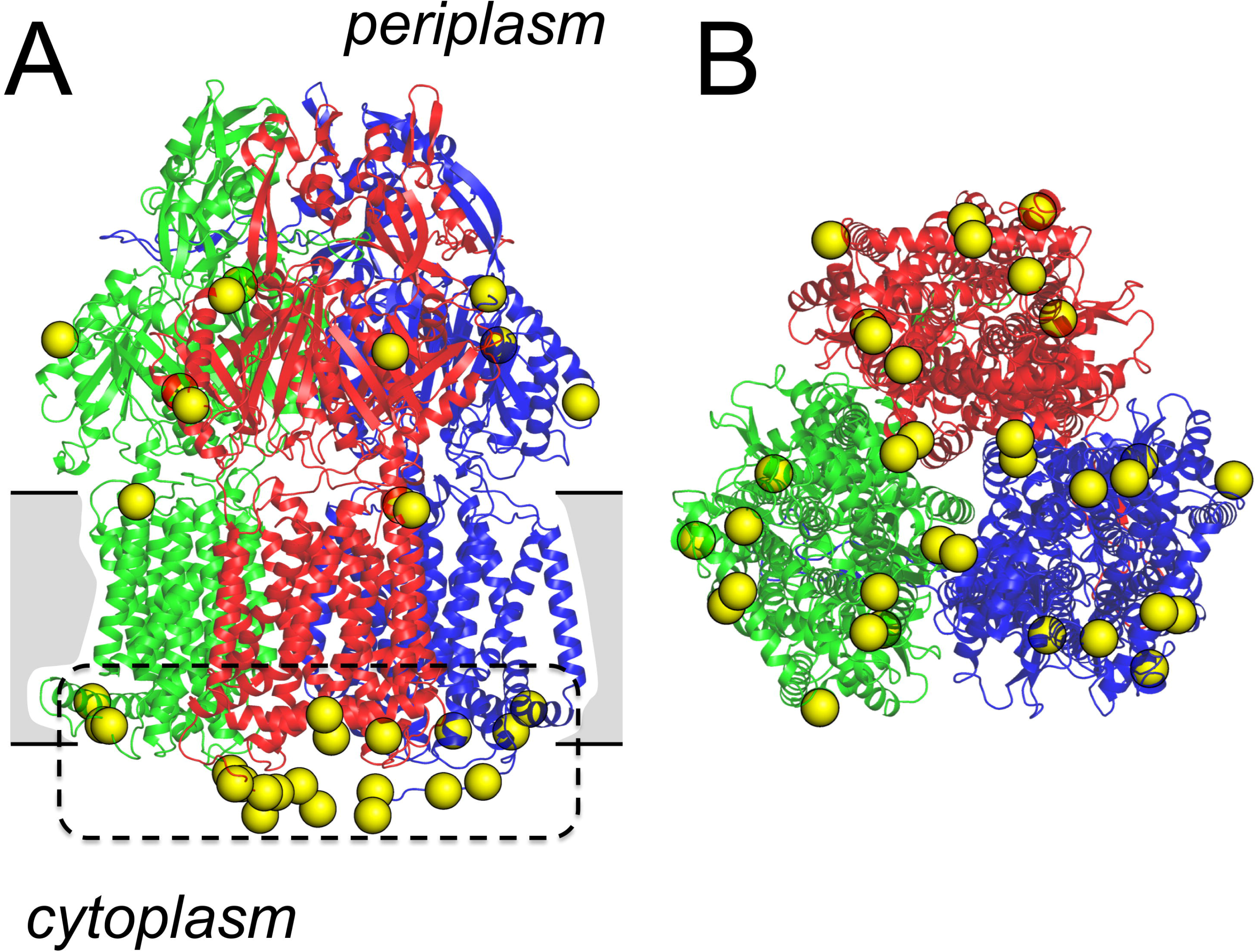
Distribution of histidine residues in the E. coli AcrB structure (PDB code: 4DX5) (23). A) View perpendicular to the membrane and B) view from the cytoplasmic side of AcrB. Chains A, B and C are coloured blue, green and red, respectively. Yellow spheres indicate the positions of histidine residues. A cluster of histidine residues on the cytoplasmic proximal face is outlined with a dashed line. Images rendered using MacPyMOL (24).

**Table 1.**
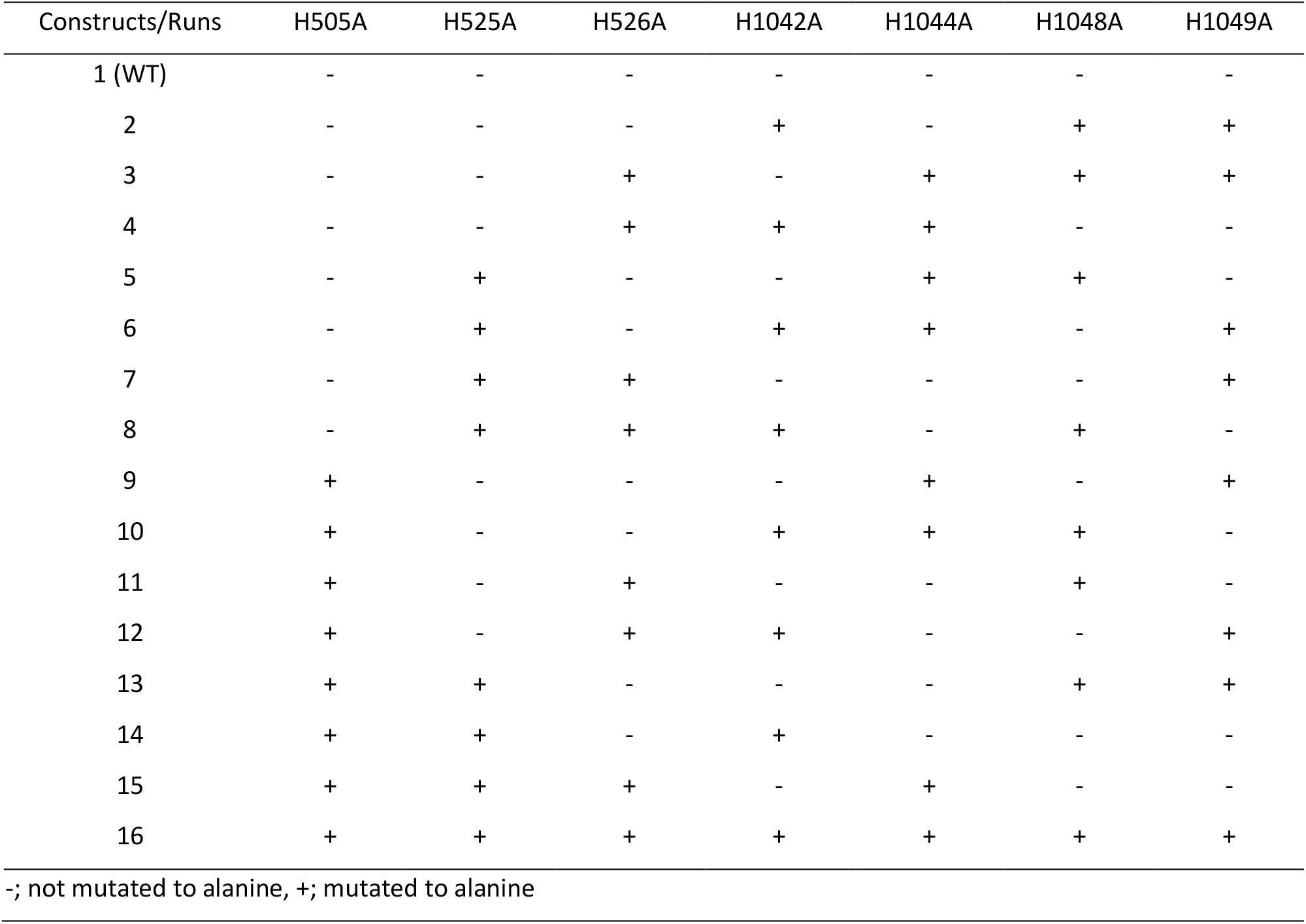
Design of the fractional factorial experiment.

| Constructs/Runs | H505A | H525A | H526A | H1042A | H1044A | H1048A | H1049A |
| --- | --- | --- | --- | --- | --- | --- | --- |
| 1 (WT) | - | - | - | - | - | - | - |
| 2 | - | - | - | + | - | + | + |
| 3 | - | - | + | - | + | + | + |
| 4 | - | - | + | + | + | - | - |
| 5 | - | + | - | - | + | + | - |
| 6 | - | + | - | + | + | - | + |
| 7 | - | + | + | - | - | - | + |
| 8 | - | + | + | + | - | + | - |
| 9 | + | - | - | - | + | - | + |
| 10 | + | - | - | + | + | + | - |
| 11 | + | - | + | - | - | + | - |
| 12 | + | - | + | + | - | - | + |
| 13 | + | + | - | - | - | + | + |
| 14 | + | + | - | + | - | - | - |
| 15 | + | + | + | - | + | - | - |
| 16 | + | + | + | + | + | + | + |

-; not mutated to alanine, +; mutated to alanine

*E. coli* AcrB with an N-terminal GFP fusion was constructed and each combination of mutations specified by the fractional factorial design was produced by site-directed mutagenesis (Table 1). Each construct was expressed in replicate in AcrB knockout *E. coli* and crudely purified on small-scale nickel affinity columns in parallel. We were unable to obtain construct 14 at the time of running the experiment, but due to the robust nature of the fractional factorial methodology, missing values can be tolerated and therefore we proceeded regardless. The effect of histidine mutants on the binding of AcrB to nickel resin could be observed by measuring in-gel GFP signals (Fig. 2 **and Fig. S1**).

**Figure 2.**
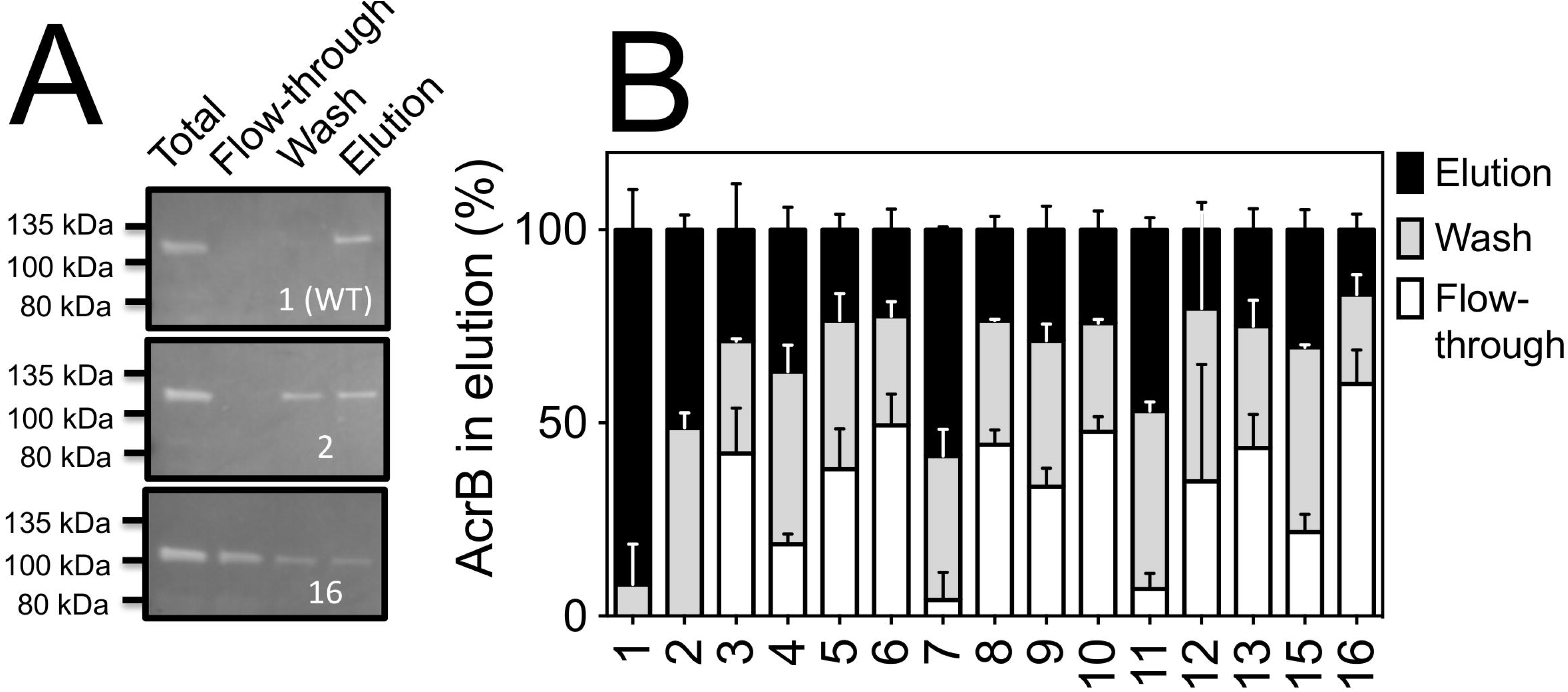
Effect of histidine mutagenesis on the ability of AcrB to bind to nickel resin. A) Example GFP fluorescence in the total, flow-through, wash and elution samples for three constructs from the fractional factorial design. B) Normalised histogram of quantified GFP signal for AcrB in the “flow-through”, “wash” and “elution” from nickel sepharose purification shows effect of mutagenesis on binding of AcrB to nickel. Error bars: standard deviation from three independent repeats.

Statistical analysis of the relative amount of GFP fluorescence in the elution allowed us to determine the main effects; we could determine which mutations to AcrB had the most significant effect on nickel binding (Table 2). Refinement of the model was carried out to include only the most significant main and two-way effects, confirming that these contributions were highly significant (Table 3).

**Table 2.** Model 1 – Equal contributions.

|  | effect | p-value | significance |
| --- | --- | --- | --- |
| (Intercept) | 9.131 |  |  |
| H505A | -1.714 | 0.1303 |  |
| H525A | -1.463 | 0.1868 |  |
| H526A | -0.685 | 0.5152 |  |
| H1042A | -1.623 | 0.1486 |  |
| H1044A | -2.314 | 0.0538 | . |
| H1048A | -1.398 | 0.2047 |  |
| H1049A | -1.015 | 0.3437 |  |

Signif. codes: 0 '\*\*\*' 0.001 '\*\*' 0.01 '\*' 0.05 '.' 0.1 ' ' 1
Residual standard error: 14.64 on 7 degrees of freedom
(1 observation deleted due to missingness)
Multiple R-squared: 0.7262, Adjusted R-squared: 0.4524
F-statistic: 2.652 on 7 and 7 DF, p-value: 0.1108

**Table 3.** Model 2 - Refined model to include most significant main and two-way effects.

|  | effect | p-value | significance |
| --- | --- | --- | --- |
| (Intercept) | 22.745 |  |  |
| H505 | -6.84 | 0.000244 | *** |
| H525 | -6.156 | 0.000465 | *** |
| H1042 | -6.592 | 0.000306 | *** |
| H1044 | -4.149 | 0.004299 | ** |
| H505:H1044 | 5.834 | 0.000641 | *** |
| H525:H1044 | 3.929 | 0.005682 | ** |
| H1042:H1044 | 5.592 | 0.000823 | *** |

Signif. codes: 0 '\*\*\*' 0.001 '\*\*' 0.01 '\*' 0.05 '.' 0.1 ' ' 1
Residual standard error: 5.365 on 7 degrees of freedom (1 observation deleted due to missingness)
Multiple R-squared: 0.9632, Adjusted R-squared: 0.9264
F-statistic: 26.18 on 7 and 7 DF, p-value: 0.0001655

The refined model (Table 3) clearly shows that mutation of H505, H525, H1042 and H1044 have the most significant effect on reducing the affinity of *E. coli* AcrB for nickel (Fig. 3). Notably, the effects of each mutation are not additive, particularly in the case of H1044, which does not give any further improvement in the presence of the other mutations, but can replace any one of them to give essentially identical effects (Table 4). This result suggests that a synergistic contribution of the histidine residues is responsible for nickel binding, agreeing with the hypothesis that several spatially close histidine residues are required for nickel ion coordination. Therefore, mutations to H505, H525 and H1042 will produce *E. coli* AcrB with low affinity to nickel, but any one of these mutations could be replaced by mutation of H1044 to get essentially the same result.

**Figure 3.**
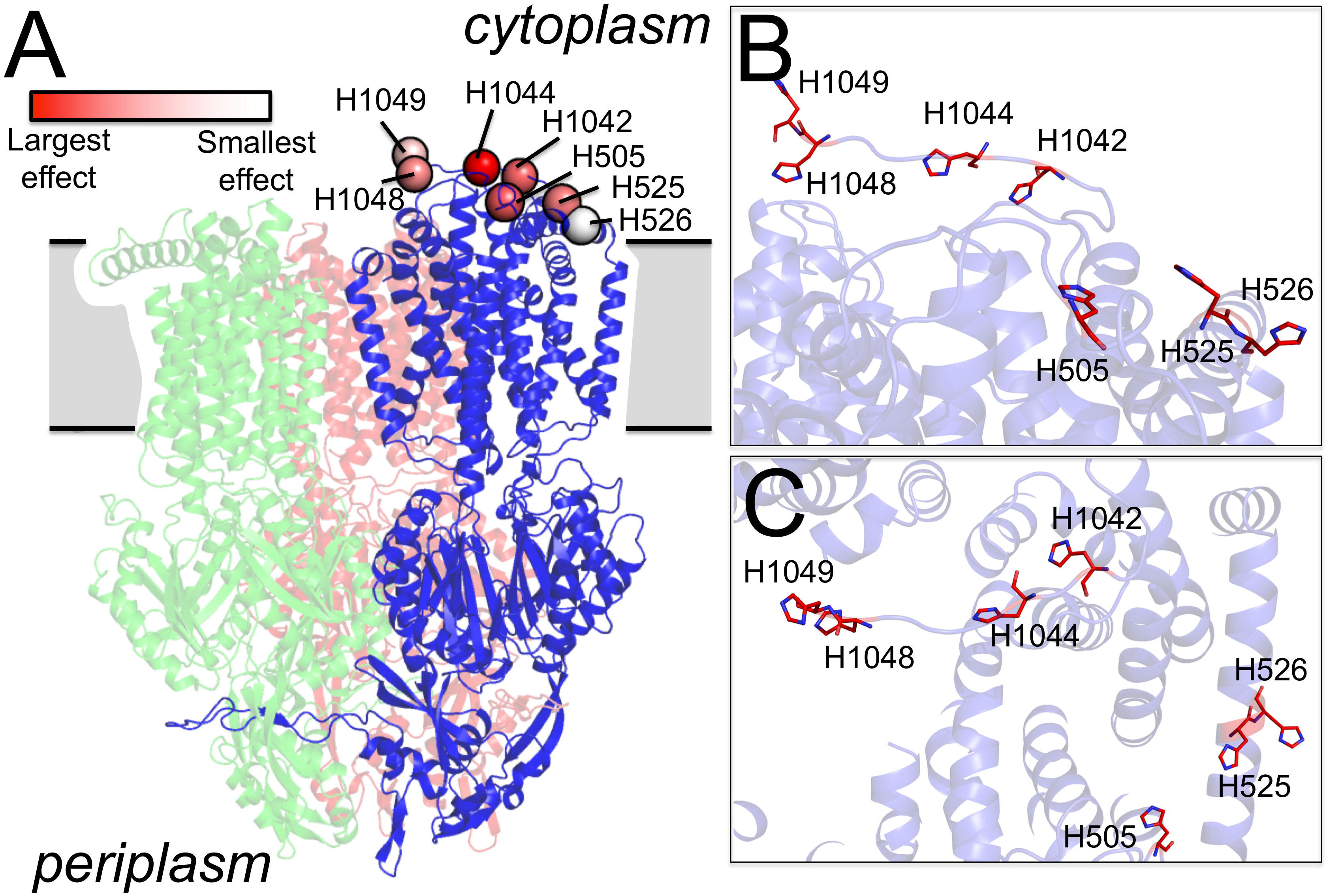
A) Cartoon representation of AcrB trimer (PDB: 4DX5). Chain is represented in bold with the positions of seven histidine residues represented as spheres. The colour of the spheres indicates the strength of the effect of their mutation on nickel resin binding (as detailed in the key). A detailed view of the histidine positions in AcrB from the side (panel B) and top (panel C). Residues are rendered as red sticks with positions of nitrogen coloured blue. Images rendered using MacPyMol (24).

**Table 4.** Two-way effects.

| H505 |  |  |  | H525 |  |  |  | H1042 |  |  |  |
| --- | --- | --- | --- | --- | --- | --- | --- | --- | --- | --- | --- |
|  |  | - | + |  |  | - | + |  |  | - | + |
| H1044 | - | 56.3 | 20.2 | H1044 | - | 52.6 | 23.9 | H1044 | - | 55.6 | 27.9 |
|  | + | 27.9 | 25.0 |  | + | 29.6 | 23.3 |  | + | 20.9 | 25.0 |

There is a caveat to add; due to the nature of the minimal design, we cannot be sure that the large interactions we see are really due to the mutations they are labelled by. For example, the interaction labelled H505:H1044, really estimates this plus H525:H526 plus H1042:H1049, but given that the main effects of H526 and H1049 are close to zero, it would be a strange system that gave this result. It would mean that, for example, H526 had a large beneficial effect in the absence of H525 and a large detrimental effect in the presence of H525 and these two effects were of almost exactly the same size.

To confirm that mutations to residues H505, H525, H1042 and H1044 could produce an AcrB construct with reduced affinity for nickel, those mutations were combined, and an extensive purification procedure was tested; washing the nickel sepharose resin with 10 column volumes of wash buffer (Fig. 4). There was significantly less (p > 0.01) AcrB eluted from nickel sepharose when residues H505, H525, H1042 and H1044 were mutated to alanine in comparison to AcrB with wild-type sequence (Fig. 4), most of the AcrB had eluted during the wash steps. This result confirms that this combination of mutations are the optimum for creating a low nickel affinity AcrB construct.

**Figure 4.**
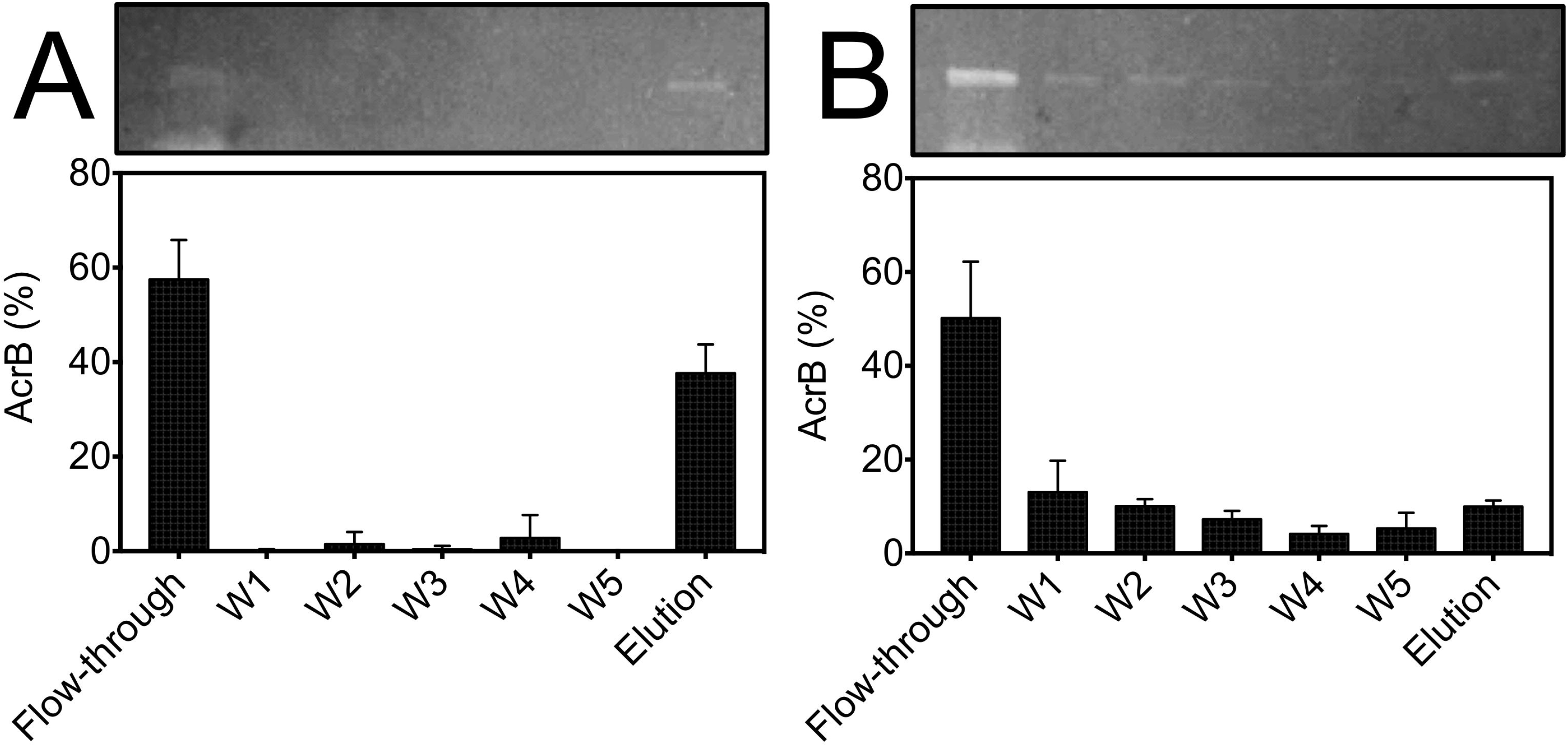
Comparison of nickel sepharose binding between A) GFP fusion with wild-type AcrB and B) GFP fusion with AcrB tetra-mutant H505A, H525A, H1042A and H1044A using 10 column volume wash (each wash step represents two column volumes). Error bars: standard deviation from three independent repeats.

## Discussion

Here, we have demonstrated the use of fractional factorial design for protein design and engineering. At the outset of the work the C-terminal residues (H1042, H1044, H1048 and H1049) were suspected to be the main contributors to nickel sepharose binding (25), but there were also histidine residues distant in sequence but spatially close to the C–terminus (H505, H525 and H526). We tested a small subset of different specific combinations of alanine replacements at these seven histidine residues in the native AcrB sequence designed in a fractional factorial screen (Table 1). Statistical analyses of the results suggested that mutations of residues H505, H525, H1042 and H1044 had the biggest effect on binding (Fig. 3), and we confirmed this to be the case experimentally (Fig. 4). This novel result is in contrast with the originally held belief that *only* C-terminal residues were important in nickel sepharose binding; the best combination of mutations could not be predicted prior to the experiment.

We note that the residues important for nickel binding form two spatially close pairs; pair H505:H525 and pair H1042:H1044 (Fig. 3), and we hypothesise that these residues are at the correct distance apart from one another to correctly coordinate the nickel ions. However, there are also spatially close pairs of histidine residues that were not indicated to be important for nickel binding, such as, H525:H526 and H1048:H1049. It is possible that histidine residues directly adjacent to one another cannot adopt the correct geometry in order to correctly coordinate nickel ions. However, this interpretation does not explain why H1044 which is ~27 Å distant from H505 and H525 in the crystal structure appears to behave in a synergistic manner with all three of the other residues indicated to be important (H505, H525 and H1042). One possibility is that any analysis based on the crystal structure alone does not account for any flexibility of the C-terminus of AcrB in solution. Indeed, of the numerous crystal structures of AcrB available in the protein data bank, the large majority of these structures are missing electron density for the C-terminal region, indicating that this is a flexible part of the protein. The position of H1044 on the end of the flexible C-terminus may allow it to come closer to H505 and H525 in solution in order to assist in the coordination of a nickel ion.

The fractional factorial design allowed us to sample a large mutational space (2^7^ combinations) with just an eighth of this total number of combinations of mutations. The fractional factorial design allowed a thorough investigation of mutations that reduce binding of AcrB to nickel sepharose, but reduced the amount of work and material costs by a factor of eight; we could understand the effect of mutating everything in every combination while only having to perform an eighth of that total experiment. Furthermore, we were able to handle the absence of results for one of the tests in the series without losing information about the main effects, highlighting one of the strengths of the fractional factorial methodology. This attribute of the fractional factorial design would be highly desirable in high-throughput cloning campaigns as is generally required for protein engineering, as absences of some mutations due to errors in cloning or expression can be ignored without significant detriment to the understanding of main-effects in the system.

There would be significant room for expansion for this technique. Here, we have chosen a system that was manageable on a small scale; however, with the use of high-throughput cloning methods as often applied for other protein engineering applications there is no reason this technique could not be expanded to cover an even larger mutational space. For example, here we have concentrated on mutating each position to just one other residue (alanine), and a third amino acid could easily be added without making the scale of the experiment too large to handle: for a full factorial of that experiment, 3^7^ combinations would be required, but using fractional factorial design the space could be sampled with just 82 combinations of mutants in a 1/27 experiment.

The specific use of fractional factorial design demonstrated here validates the use of this method for protein engineering, and provides a framework to apply it broadly for many other applications. For example, we believe this could have important application in the investigation of altering enzyme active site residues to change affinity for substrate or alter substrate preference. In the case of active site residues, it is often clear which residues form the most important interactions with substrate to define specificity or catalytic activity, but unclear what combination of changes to those residues (of the 20 amino acids) will have the desired effect on enzyme catalysis. We propose that fractional factorial design would provide an excellent framework to allow comprehensive understanding of the effect of changing all residues in an active site in all combinations, allowing the sampling of a broad range of possible ways to modify the properties of the enzymatic reaction.

We also see a broad benefit of using fractional factorial design for altering the residues of antibody complementarity determining regions (CDR) in order to improve the affinity of the antibody for its epitope. Typically, antibody maturation and CDR improvement is done using random mutagenesis. However, as discussed above, there are biases in random mutagenesis that will prevent the full range of mutational space from being accessed. We propose that a fractional factorial approach would allow a much broader sampling of the possible mutational space, and by limiting mutation to just the CDRs the experiments will not be unfeasibly large.

In the case of protein stabilisation, fractional factorial design may not be able to replace scanning or random mutagenesis methods for the initial identification of single positions with beneficial effects to protein stability due to the staggering large number of possible combinations even in a small protein. However, fractional factorial design can be extremely valuable to help determine which of the mutations initially identified by other methods should be combined, and suggesting the minimal number of changes required for maximal effect.

In combination with stability assays, we also envisage the use of fractional factorial design to infer two-way effects (pairs of residues that do not have an additive effect) allowing us to experimentally determine the proximity of residues to one another. This type of information can be highly informative in proteins of unknown structure, as these residue pairs can act as distance constraints for guiding and improving computationally derived protein models.

## Materials and methods

### Fractional factorial design

The *E. coli* AcrB residues H505, H525, H526, H1042, H1044, H1048 and H1049 were taken as the seven factors for investigation, with two levels for each factor to be investigated (non-mutated; - or mutated to alanine; +) (Table 5)

**Table 5.**
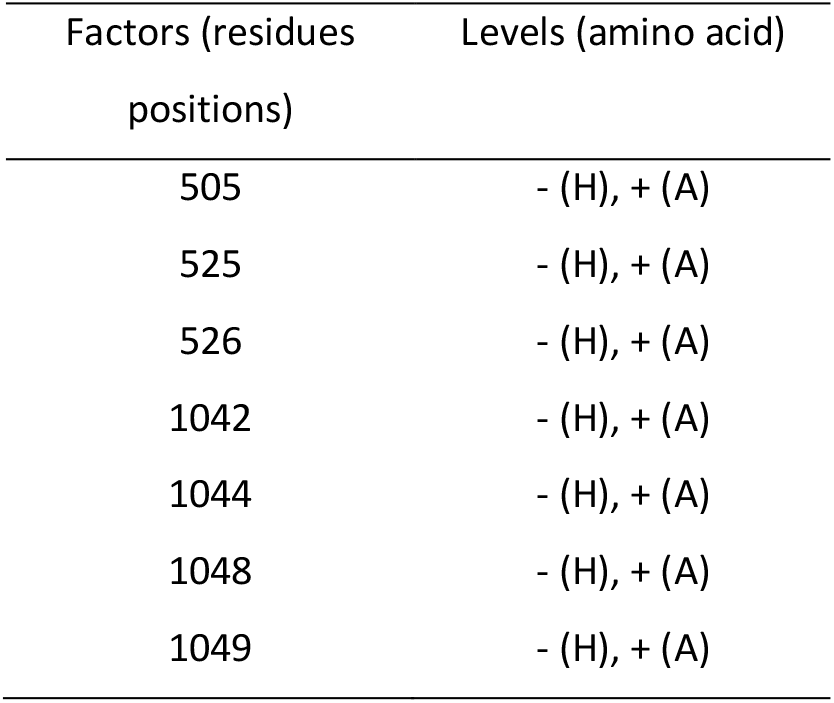

A 1/8 design was used (16 runs in the fractional factorial design vs 128 runs in the full factorial design) (Table 1), which can provide information about main effects and some two-way effects can be inferred.

### AcrB mutagenesis

The *E. coli* AcrB gene was cloned into a pET-21-GFP vector (pET-21-GFP-AcrB) to create the initial GFP-tagged AcrB construct. Mutagenic primers were designed using either “QuickChange” or “Round-the-Horn” methods (26). Mutations were introduced into AcrB sequentially as constructs required between three and seven mutations in total. Briefly, 10 µL PCR reactions were setup using mutagenic primers, Q5 DNA polymerase (NEB, Ipswich, USA) and the pET21-GFP-AcrB template (at approximately 10 ng/µL). The reaction was carried out (Thermal Cycler, Bio-Rad, Hercules, USA) with primer annealing temperatures determined theoretically, and a long elongation time (30 seconds per kbp; 3.5 minutes). Following PCR the reactions were treated with either DpnI or a mixture of T4 DNA Ligase, T4 Polynucleotide Kinase and DpnI for the QuickChange or “Round-the-Horn” methods, respectively (all enzymes were supplied by NEB, Ipswich, USA). These reactions were incubated at room temperature for 1 hour before transformation into chemically competent OmniMAX *E. coli* cells, plating on LB agar plates containing 100 µg/mL carbenicillin and overnight incubation at 37°C. Correctly mutated plasmids were confirmed by sanger sequencing (Eurofins genomics, Luxembourg, Switzerland) after mini-prep plasmid purification (Nucleospin Plasmid kit; Macherey-Nagel, Düren, Germany) from overnight culture of single colonies in LB containing 100 µg/mL ampicillin and incubation at 37°C.

### AcrB expression and quantification of affinity for nickel

Chemically competent *E. coli* strain C41 *ΔAcrB* pRARE2 were transformed with the 16 pET-21-GFP-AcrB constructs using heat-shock method and plated onto LB agar plates containing 100 µg/mL carbenicillin. Three single colonies were selected for each AcrB construct and used to inoculate 4 mL of auto-induction media (Na2HPO4, 10 mM, KH2PO4, 5 mM, tryptone, 0.2 % (w/v), yeast extract, 0.05 % (w/v), NaCl, 20 mM, Glycerol, 0.6 % (v/v), glucose, 0.05 % (w/v), lactose 0.2 % (w/v), 100 µg/mL ampicillin). Cultures were grown in sterile 24-well deep-well blocks, incubated at 30°C with shaking at 200 rpm for 6 hours before cooling to 18°C with shaking at 200 rpm overnight.

Bacterial pellets were collected by centrifugation at 3000 x g for 10 mins and washed once with 1 mL MilliQ-H2O. Pellets were re-suspended in 250 µL of Lysis/Solubilisation buffer (20 mM Tris-HCl pH 8, 300 mM NaCl, 1.5 % (w/v) dodecyl maltoside, 2 µg/mL DNAse I, 3 x EDTA-free protease inhibitor cocktail) and solubilised with mixing at 4°C for 1 hour. 100 µL of pre-equillibrated nickel sepharose resin (Ni sepharose 6 ff, GE Healthcare, Chicago, USA) slurry was transferred to each well of a UniFilter GF/B pore size 1 µm Conical Bottom 96-well Filter Plate (Whatman, Little Chalfont, UK) and spun dry (1000 x g, 5 mins). Solubilised cell lysate was centrifuged for 10 minutes at 3000 x g, 4°C before 200 µL of each condition was applied to the 96-well filter plate and protein was allowed to batch-bind with the nickel resin at 4°C at 1000 rpm (Eppendorf MixMate; Eppendorf, Hamburg, Germany).

After batch binding, the plate was centrifuged at 1000 x g for 1 min and “flow-through” collected. 200 µL of Buffer A (20 mM Tris-HCl pH 8, 300 mM NaCl, 50 mM imidazole, 0.05 % (w/v) dodecyl maltoside) was then added to each well and incubated for a further 10 mins at 4°C with mixing at 1000 rpm (Eppendorf MixMate; Eppendorf, Hamburg, Germany). The plate was centrifuged at 1000 x g for 1 min and “wash” collected. 200 µL of Buffer B (20 mM Tris-HCl pH 8, 300 mM NaCl, 250 mM imidazole, 0.05 % (w/v) dodecyl maltoside) was then added to each well and incubated for a further 10 mins at 4°C with mixing at 1000 rpm (Eppendorf MixMate; Eppendorf, Hamburg, Germany). The plate was centrifuged for a final time at 1000 x g for 1 min and “elution” collected. 50 µL samples from the “solubilised lysate”, “flow-through”, “wash” and “elution” were mixed with 5 x SDS-PAGE loading buffer (250 mM Tris-HCl, pH 6.8, 10 % SDS, 30 % (v/v) glycerol, 10 mM DTT, 0.05 % (w/v) Bromophenol Blue). Samples were loaded onto 15-well 4-20 % Mini- PROTEAN pre-cast PAGE gels (Bio-Rad, Hercules, USA) and run for 1 hour at 150 V in SDS-PAGE running buffer (25 mM tris-HCl pH 8.3, 193 mM glycine, 0.1 % (w/v) SDS). In-gel GFP fluorescence was visualised (G:BOX Chemi XX6 with Blue LEDs; Syngene, Gurgaon, India) before Coomassie staining (QuickStain; Generon, Slough, UK). In-gel GFP fluorescence was analysed using Fiji (27) to determine the pixel density of each band containing GFP-AcrB. The pixel density corresponding to GFP fluorescence in the elution relative to the total GFP signal across the flow-through, wash and elution was used to compare the relative affinity for nickel between the different constructs. Error bars are representative of the standard deviation over three individual repeats for each sample.

### Analysis of fractional factorial experiment

Statistical analysis of the fractional factorial experiment was carried out using R, a language and environment for statistical computing (https://www.R-project.org).

## Acknowledgements

We wish to thank Clair Philips for assisting with mutagenesis.

The *E. coli* strain C41 *ΔAcrB* pRARE2 was a kind gift from Dr. Vincent Postis

## Contributions

S. P. D. H., S. G. G. and A. G. designed the experiments. S. P. D. H, D. W., J. M and A. M. performed the mutagenesis and binding experiments. S. P. D. H., S. G. G. and A. G. analysed the data and wrote the paper.

## References

1. Cunningham B, Wells J (1989) High-resolution epitope mapping of hGH-receptor interactions by alanine-scanning mutagenesis. Science (80-) 244(4908):1081–1085.

2. Magnani F, et al. (2016) A mutagenesis and screening strategy to generate optimally thermostabilized membrane proteins for structural studies. Nat Protoc 11(8):1554–1571.

3. Magnani F, Shibata Y, Serrano-Vega MJ, Tate CG (2008) Co-evolving stability and conformational homogeneity of the human adenosine A2a receptor. Proc Natl Acad Sci 105(31):10744–10749.

4. Sarkisyan KS, et al. (2016) Local fitness landscape of the green fluorescent protein. Nature 533(7603):1–11.

5. Klenk C, Ehrenmann J, Schütz M, Plückthun A (2016) A generic selection system for improved expression and thermostability of G protein-coupled receptors by directed evolution. Sci Rep 6(November 2015):21294.

6. Tate CG (2015) Identifying Thermostabilizing Mutations in Membrane Proteins by Bioinformatics. Biophys J 109(7):1307–1308.

7. Bhattacharya S, Lee S, Grisshammer R, Tate CG, Vaidehi N (2014) Rapid Computational Prediction of Thermostabilizing Mutations for G Protein-Coupled Receptors. J Chem Theory Comput 10(11):5149–5160.

8. Yasuda S, et al. (2016) Identification of Thermostabilizing Mutations for Membrane Proteins: Rapid Method Based on Statistical Thermodynamics. J Phys Chem B 120(16):3833–3843.

9. Heydenreich FM, Vuckovic Z, Matkovic M, Veprintsev DB (2015) Stabilization of G protein- coupled receptors by point mutations. Front Pharmacol 6(MAR):1–15.

10. Wilson DS, Keefe AD (2001) Random Mutagenesis by PCR. Current Protocols in Molecular Biology (John Wiley & Sons, Inc., Hoboken, NJ, USA), p Unit8.3.

11. Echols H, Lu C, Burgers PM (1983) Mutator strains of Escherichia coli, mutD and dnaQ, with defective exonucleolytic editing by DNA polymerase III holoenzyme. Proc Natl Acad Sci 80(8):2189–2192.

12. Lee CMY, Iorno N, Sierro F, Christ D (2007) Selection of human antibody fragments by phage display. Nat Protoc 2(11):3001–3008.

13. Sauer DB, Karpowich NK, Song JM, Wang D-N (2015) Rapid Bioinformatic Identification of Thermostabilizing Mutations. Biophys J 109(7):1420–1428.

14. Carter CW, Carter CW (1979) Protein Crystallization Using Incomplete Factorial- Experiments. J Biol Chem 254(23):2219–2223.

15. Jancarik J, Kim SH (1991) Sparse matrix sampling: a screening method for crystallization of proteins. J Appl Crystallogr 24(4):409–411.

16. Papaneophytou CP, Kontopidis G (2014) Statistical approaches to maximize recombinant protein expression in Escherichia coli: A general review. Protein Expr Purif 94:22–32.

17. He GQ, Kong Q, Ding LX (2004) Response surface methodology for optimizing the fermentation medium of Clostridium butyricum. Lett Appl Microbiol 39(4):363–368.

18. Veesler D, Blangy S, Cambillau C, Sciara G (2008) There is a baby in the bath water: AcrB contamination is a major problem in membrane-protein crystallization. Acta Crystallogr Sect F Struct Biol Cryst Commun 64(10):880–885.

19. Glover CAP, et al. (2011) AcrB contamination in 2-D crystallization of membrane proteins: Lessons from a sodium channel and a putative monovalent cation/proton antiporter. J Struct Biol 176(3):419–424.

20. Psakis G, Polaczek J, Essen L-O (2009) AcrB et al.: Obstinate contaminants in a picogram scale. One more bottleneck in the membrane protein structure pipeline. J Struct Biol 166(1):107–111.

21. Eaves DJ, Ricci V, Piddock LJV (2004) Expression of acrB, acrF, acrD, marA, and soxS in Salmonella enterica serovar Typhimurium: role in multiple antibiotic resistance. Antimicrob Agents Chemother 48(4):1145–50.

22. Padilla E, et al. (2010) Klebsiella pneumoniae AcrAB efflux pump contributes to antimicrobial resistance and virulence. Antimicrob Agents Chemother 54(1):177–83.

23. Eicher T, et al. (2012) Transport of drugs by the multidrug transporter AcrB involves an access and a deep binding pocket that are separated by a switch-loop. Proc Natl Acad Sci 109(15):5687–5692.

24. Schrödinger, LLC (2015) The {PyMOL} Molecular Graphics System, Version~1.8.

25. Wiseman B, et al. (2014) Stubborn Contaminants: Influence of Detergents on the Purity of the Multidrug ABC Transporter BmrA. PLoS One 9(12):e114864.

26. Hemsley A, Arnheim N, Toney MD, Cortopassi G, Galas DJ (1989) A simple method for site- directed mutagenesis using the polymerase chain reaction. Nucleic Acids Res 17(16):6545–51.

27. Schindelin J, et al. (2012) Fiji: an open-source platform for biological-image analysis. Nat Methods 9(7):676–682.

